# Small protein modules dictate prophage fates during polylysogeny

**DOI:** 10.1101/2022.09.16.508337

**Authors:** Justin E. Silpe, Olivia P. Duddy, Fatima A. Hussain, Kevin J. Forsberg, Bonnie L. Bassler

## Abstract

DNA-damaging agents are the pervasive inducers of temperate phages in model bacteria. However, most bacteria in the biosphere are predicted to carry multiple prophages, a state called polylysogeny, making it unclear how co-residing prophages compete for host cell resources if they all respond to the identical trigger. We discover regulatory modules encoded on phage genomes that control prophage induction independently of the DNA damage cue. Genes specifying these pathways exist in linear plasmid-like phages at sites essential for phage propagation. The modules lack sequence similarity but display a shared regulatory logic of a transcription factor that activates expression of a neighboring gene encoding a small protein. The small protein inactivates the master repressor of lysis, leading to prophage induction. In some phages, the regulatory unit detects sensory information including quorum-sensing autoinducers, making lysis host-cell-density dependent. Exposure of the polylysogens studied here to different induction scenarios reveals that mixed phage populations emerge following DNA damage, however, induction through the SOS-independent module drives near-exclusive production of the phage sensitive to that specific cue. Considering the lack of potent DNA-damaging agents in natural habitats, we propose that additional phage-encoded sensory pathways that drive lysis play fundamental roles in phage-host biology and inter-prophage competition.

## INTRODUCTION

Phages are viruses that infect bacteria. Two mechanisms control phage reproduction. First, lytic phages infect the bacterial host, replicate, and kill the host cell. Second, temperate, or lysogenic, phages remain dormant in host cells and are passed down to progeny. Temperate phages can transition from the lysogenic state to the lytic state in response to DNA-damaging agents that activate the bacterial SOS response.^1–3^ During lysogeny, model temperate phages, including phage lambda, produce a repressor (called cI) that binds to and prevents expression from a promoter (termed P_R_) controlling the lysis genes.^3^ Following DNA damage, activation of the host RecA protein leads to autoproteolysis and inactivation of the cI repressor.^1,4^ Consequently, P_R_ is de-repressed, triggering phage replication, host-cell lysis, and transmission of the phage to neighboring cells. Other temperate phages harbor repressors lacking the peptidase domain responsible for autoproteolysis. Rather, a peptidase or antirepressor encoded elsewhere in the phage genome is activated by the host SOS response.^5,6^ The understanding that all bacteria possess *recA* coupled with the fact that phages are omnipresent, has led to the common view that the host SOS response is the ‘universal’ prophage inducer. However, significant concentrations of potent DNA-damaging agents are rare in the environment, and increasingly, phages are being discovered that are not induced by DNA damage.^7^ Together, these findings suggest that undiscovered induction triggers exist in nature.

Recent discoveries reveal that some phages respond to quorum-sensing (QS) signals as inputs into their lysis-lysogeny lifestyle transitions.^8–10^ QS is a process of cell-to-cell communication that bacteria use to orchestrate collective behaviors. QS relies on the production, release, and group-wide detection of and response to extracellular signaling molecules called autoinducers (AI).^11^ Phages can harbor phage-phage QS-like communication systems, such as the arbitrium system in SPβ phages,^10^ or as in vibriophage VP882, they can monitor host bacterial QS-mediated communication pathways to tune the timing of the lysis-lysogeny switch to changes in host-cell density.^9^ Phage VP882 is a linear plasmid-like prophage that encodes a homolog of the *Vibrio* QS receptor VqmA, called VqmA_Phage_, which is activated by a host-produced AI, called DPO.^9,12^ Upon binding DPO, VqmA_Phage_ activates transcription of a counter-oriented gene called *qtip*. Production of Qtip launches host-cell lysis. Our hypothesis is that, by surveilling a bacterial-produced QS signal, the phage can factor host-cell density information into its decision-making process to maximize the probability of infecting other cells in the population.

In this work, we uncover phage-encoded regulatory components that operate in addition to, and independently of, the canonical SOS response. These SOS-independent programs were discovered by systematically identifying linear plasmid-like phages in existing phage genome databases and selecting only those containing autoproteolytic cI repressors. This subset of linear phages, which are SOS-responsive by virtue of the repressor, were subjected to a genome neighborhood-based search for the presence of other genes possibly encoding lysis regulatory components in conserved phage loci (near genes encoding linearization, replication, and partition machinery). Prophages that could be matched to an obtainable lysogen were prioritized for study. Using this approach, we discover multiple phage-encoded lysis-inducing modules that, despite bearing little resemblance to one another at the sequence level, share a common regulatory logic. They all employ a transcription factor to activate the expression of a divergently transcribed gene encoding a small protein (smORF). The smORF launches the transition from lysogeny to lysis. The smORFs studied here lack characterized homologs and predictable domains, and yet they operate by inactivating the same respective target, the cI repressor in the phage that encodes it. The mechanism of regulation of the smORF by its partner transcription factor can vary. In some cases, the transcription factor operates independently to activate *smORF* gene expression. In other cases, *smORF* gene expression requires a xenobiotic responsive element (XRE) family protein working in conjunction with a LuxR-type QS receptor/transcription factor that requires a bacterial-produced AI ligand for activity. Finally, most of the prophage-containing isolates we study are polylysogenic. We show that the addition of a DNA-damaging agent to these polylysogens leads to phage-mediated lysis of the bacteria and release of a mixed population of phage particles. Unlike DNA damage, induction via the newly discovered regulatory module results in near-exclusive production of the phage responsive to the specific input. Our results suggest that activities of these SOS-independent pathways dictate the outcomes of prophage-prophage competition by expanding the range of stimuli to which specific prophages can respond.

## RESULTS

### A bioinformatic search for SOS-inducible linear plasmid-like phages reveals phages that encode additional lysis-lysogeny regulatory modules

In vibriophage VP882, the genes *vqmA_Phage_* and *qtip* reside between *repA* and *telN*, hallmarks of all known linear plasmid-like phages. Inspired by this arrangement, we conducted a search among sequenced phages for putative novel lysis-lysogeny modules located between *repA* and *telN*. In addition to NCBI, we searched six recently curated phage and phage-plasmid databases^13–18^ spanning diverse environmental, marine, and human body sites for convergently-oriented *repA* and *telN* genes residing within 10 kilobases of each other. The search revealed 784 putative linear plasmid-like phage genomes, 274 of which contained unique *repA-telN* loci (Supplementary Table 1). In 271 of 274 genomes (99%), the gene upstream of *repA* encoded a cI-like DNA-binding protein that was adjacent to putative, divergently transcribed lytic, structural, and regulatory genes. A panel of representative loci is shown in Figure 1a. We focused on phage genomes harboring genes encoding RecA-dependent, autoproteolytic cI repressors under the assumption that any additional regulatory modules we uncover would likely respond to SOS-independent inputs. Filtering phage genomes using these criteria led to 61 unique loci. The majority of phages eliminated at this step (210/271) encoded *repA*-*telN* loci resembling those of the *Escherichia coli* linear plasmid-like phage called N15. This finding is likely a consequence of the overrepresentation of *Enterobacteriaceae* in the sequencing databases. N15 is known to be subject to antirepression,^19,20^ and we determined the repressor it encodes is non-proteolytic (Figure 1b). In contrast, phages with putative autoproteolytic repressors were less prevalent and more diverse in these databases. For instance, when we clustered *repA*-*telN* loci at 80% identity, there were fewer sequences per cluster for those with autoproteolytic repressors than non-proteolytic repressors, indicating that our autoproteolytic set less redundantly sampled nature’s diversity (p = 0.016, Student’s t-test, Supplementary Table 2). Moreover, despite there being three times more nonproteolytic than autoproteolytic phages present in our sample, the number of singleton clusters formed of the autoproteolytic phages (29) was higher than that for the non-proteolytic phages (26) (Supplementary Table 2). Highlighting their understudy, the only member of the identified set of phages with autoproteolytic repressors that has been investigated in this regard is phage VP882 (Figure 1a and 1b), described above, and a singleton in our current cluster analysis. Many of the phages from our database searches come from metagenomic sequencing projects, are unobtainable, and/or have no known host. Nonetheless, with the goal of probing the functions of these putative sensory input pathways, we focused on obtainable phage-host pairs.

**Figure 1.**
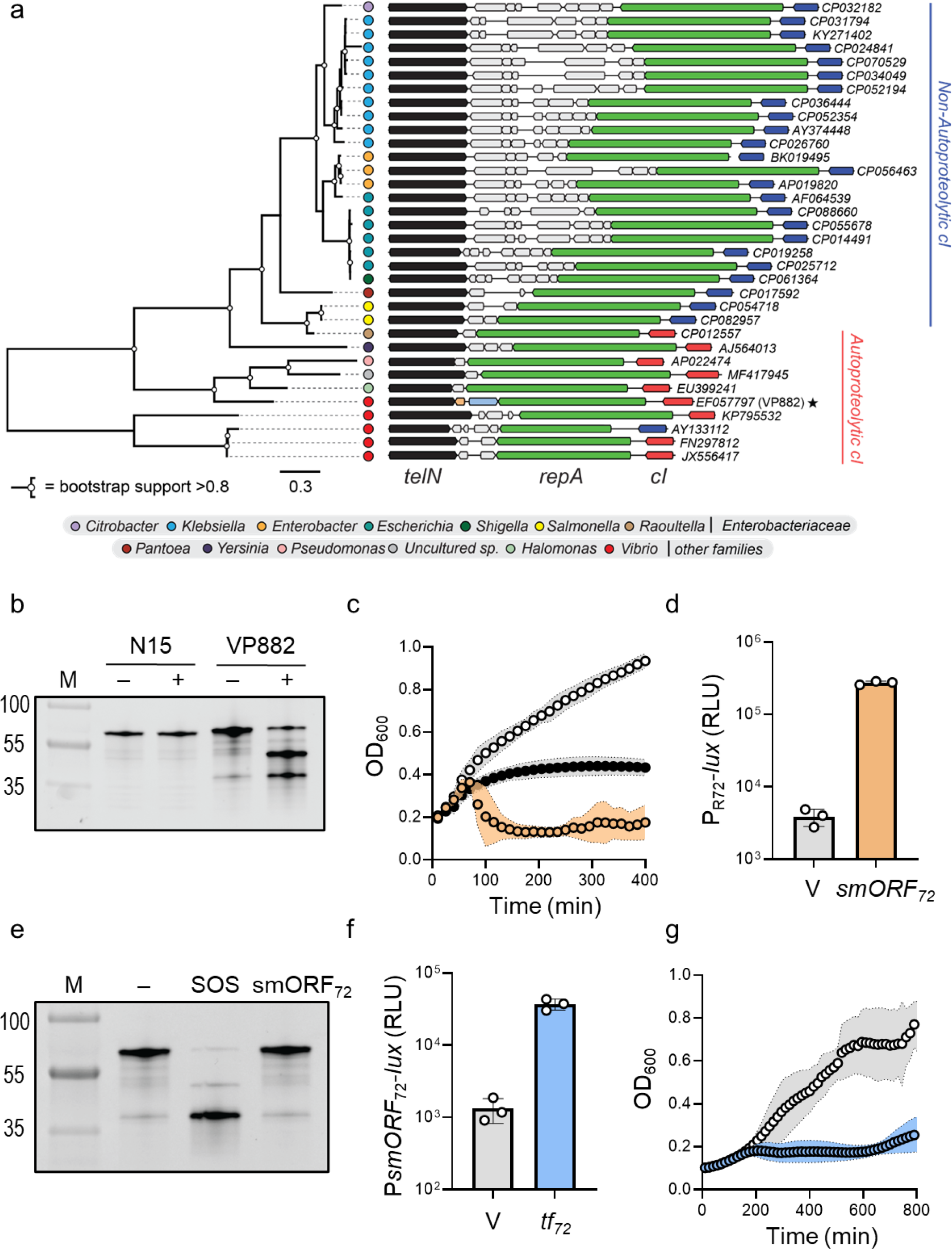
Variable gene content in an otherwise conserved locus of linear plasmid-like phages reveals TF-smORF modules that regulate lysis independently of SOS. **(a)** Phylogenetic tree (left) of 34 representative TelN proteins. TelN protein groups typically segregate with bacterial genera (colored circles, legend at bottom). Nodes with bootstrap support >0.8 are indicated with white circles. Gene neighborhoods (right) for the 34 loci encoding convergently-oriented *telN* (black) and *repA* (green) genes. The variable loci encoding *tf-smORF* modules are located between *telN* and *repA*. Phage VP882 is denoted with a star and the SOS-independent pathway components (*vqmA_Phage_* and *qtip*) are colored blue and orange, respectively. All other genes are colored gray, encode unknown functions, and vary across loci. Genes encoding predicted autoproteolytic repressors (red) cluster together with respect to TelN phylogeny and are distinct from genes encoding non-cleavable repressors (navy). NCBI accession numbers are depicted to the right of each sequence. The scale bar indicates the number of amino acid substitutions per site. **(b)** SDS-PAGE in-gel labeling of the non-proteolytic N15 phage repressor (HALO-cI_N15_) and the autoproteolytic phage VP882 repressor (HALO-cI_VP882_). − and + indicate, respectively, the absence and presence of 500 ng mL^−1^ ciprofloxacin used to induce the SOS response. M denotes the molecular weight marker (representative bands are labeled). **(c)** Growth of *Vibrio* 1F-97 carrying aTc-inducible *smORF_72_* in medium containing 50 ng mL^−1^ aTc (orange), 500 ng mL^−1^ ciprofloxacin (black), or water (white). **(d)** P_R72_-*lux* expression in *E. coli* carrying an empty vector (designated V) or aTc-inducible *smORF_72_* grown in medium containing aTc. The P_R72_-*lux* plasmid carries *cI_72_*, which natively represses reporter expression. Relative light units (RLU) were calculated by dividing bioluminescence by OD_600_. aTc concentration as in (c). **(e)** SDS-PAGE in-gel labeling of the *Vibrio* 1F-97 phage 72 repressor (cI_72_-HALO) produced by *E. coli* carrying aTc-inducible *smORF_72_*. The treatments -, SOS, and smORF_72_ refer to water, ciprofloxacin, and aTc, respectively. M as in (b). aTc and ciprofloxacin concentrations as in (c). **(f)** P*smORF_72_*-*lux* expression from *E. coli* carrying an empty vector (V) or aTc-inducible *tf_72_* grown in medium containing aTc. RLU as in (d). aTc concentration as in (c). **(g)** Growth of *Vibrio* 1F-97 carrying aTc-inducible *tf_72_* in medium lacking or containing aTc (white and blue, respectively). aTc concentration as in (c). Data are represented as means ± std with *n*=3 biological replicates (c, d, f, g).

### Transcription factor-small ORF modules regulate prophage induction independently of SOS

The first isolate we investigated is *Vibrio cyclitrophicus* 1F-97 (here forward called *Vibrio* 1F-97), which encodes a putative phage on contig 72, hereafter called phage 72. Between the phage 72 *repA* and *telN* genes are genes encoding a putative transcription factor and a counter-oriented small, 171 nt ORF (hereafter TF_72_ and smORF_72_, respectively). The sequences of these genes lack nucleotide-level identity to the *vqmA_Phage_* and *qtip* genes, however, their arrangement parallels that for *vqmA_Phage_* and *qtip* in phage VP882. Our first goal was to determine if the region between *repA* and *telN* on phage 72 controls the phage lysis-lysogeny transition. To test this possibility, we cloned *smORF_72_* under the control of an anhydrotetracycline (aTc)-inducible promoter on a plasmid (pTet-*smORF_72_*) and conjugated it into *Vibrio* 1F-97. Addition of aTc to this recombinant strain led to a precipitous decline in OD_600_ similar to that of the potent SOS-activator ciprofloxacin, indicating phage-driven lysis occurred (Figure 1c). aTc administered to *Vibrio* 1F-97 harboring an empty vector or to *E. coli* harboring the pTet-s*mORF_72_* vector did not affect growth (Extended Data Figure 1a and 1b, respectively). Thus, lysis required the presence of the phage and induction of the *smORF_72_* gene. Phage preparations obtained from *Vibrio* 1F-97 treated with aTc or ciprofloxacin contained phage 72 particles showing that phage 72 can be induced by both SOS-independent and SOS-dependent pathways (Extended Data Figure 1c).

We next explored the mechanism by which induction of smORF_72_ promotes lysis. Important for this step is that our bioinformatic analysis revealed that 99% of the *repA-telN* loci (271/274) contain genes upstream of *repA* that encode predicted phage repressor proteins (called cI_72_ and its target promoter, P_R72,_ respectively, for phage 72). We verified the function of this repressor-promoter pair by fusing the P_R72_ promoter to *lux* on a plasmid (P_R72_-*lux*). Recombinant *E. coli* carrying the construct made light, which decreased 500-fold when the gene encoding cI_72_ was also introduced (Extended Data Figure 1d). We predicted that smORF_72_ could induce lysis via inactivation of the cI_72_ repressor. Indeed, introduction of pTet-*smORF_72_* into *E. coli* carrying the *cI_72_*-P_R72_-*lux* plasmid restored high levels of light production (Figure 1d). Unlike DNA damage, smORF_72_ did not lead to cI_72_ proteolysis (Figure 1e). We conclude that smORF_72_ is an antirepressor that acts directly but nonproteolytically to inactivate its partner cI_72_ repressor protein.

We wondered how *smORF_72_* expression is naturally regulated in phage 72. As noted, a gene encoding a transcription factor, *tf_72_* is adjacent to and divergently transcribed from *smORF_72_*, providing us a logical candidate. We fused the promoter of *smORF_72_* to *lux* (P*smORF*_72_*-lux*) and transformed this reporter into recombinant *E. coli* harboring aTc-inducible *tf_72_* or an empty vector control. Induction of TF_72_ production increased light output by 28-fold compared to the empty vector (Figure 1f). Likewise, production of TF_72_ in *Vibrio* 1F-97 led to an increase in *smORF_72_* transcription (90 min after TF_72_ induction, Extended Data Figure 1e), after which the culture lysed (Figure 1g). These results demonstrate that TF_72_, via transcriptional activation of *smORF_72_*, drives phage 72-mediated lysis.

### Vibrio *1F-97 harbors at least two plasmid-like phages that control lysis via transcription factor-smORF modules encoded in different genomic regions*

To survey the distribution of smORF_72_-TF_72_-like phage regulatory modules, we performed bioinformatic searches (BLASTp) of NCBI for each component. BLASTp of smORF_72_ returned no hits. Unexpectedly, BLASTp using TF_72_ as the query revealed that the gene encoding the closest homolog (~53% amino acid pairwise identity) in the NCBI database resides on a different contig within the same *Vibrio* 1F-97 genome, contig 63. Contig 63, like phage 72, harbors signatures of a linear plasmid-like phage (e.g., genes resembling *telN* and *repA*; Figure 2a), but it did not appear in our initial search because it was not in the NCBI nt database at that time (see Methods). We determined that the replication machinery encoded on contig 63, like that of phage 72, was functional by ligating an antibiotic resistance cassette to each phage *repA* gene and demonstrating antibiotic-selective growth in *E. coli*. Furthermore, DNA corresponding to contig 63 was present in phage preparations of ciprofloxacin-treated *Vibrio* 1F-97 cultures (Extended Data Figure 2a). We now refer to this element as phage 63. Phage 72 and phage 63 share little identity on a whole-genome basis (Mauve), and unlike phage 72, cloning and expression of the phage 63 DNA intervening *repA* and *telN* did not induce lysis (Extended Data Figure 2b and 2c). Rather, the gene encoding the homologous transcription factor in phage 63 (*tf_63_*) is located between genes encoding predicted partition machinery (*parAB*) and an operon encoding predicted late genes (tail and assembly genes; Figure 2a). Intriguingly, *tf_63_* resides adjacent to but in the opposite orientation from a gene encoding another hypothetical 183 nt smORF (*smORF_63_*). We hypothesized that, in phage 63, the *par*-associated, *tf_63_-smORF_63_* pair may encode proteins that perform analogous functions as the *tf_72_*-*smORF_72_* pair in phage 72 and possibly other *telN-repA* associated pairs (e.g., *vqmA_Phage_-qtip* in phage VP882). To test this possibility, we constructed the analogous set of tools described above for phage 72. In *E. coli*, production of TF_63_ led to a 50-fold increase in light production from a P*smORF_63_-lux* reporter (Figure 2b). Moreover, smORF_63_ production relieved cI_63_-mediated repression of a P_R63_*-lux* reporter and the cI_63_ repressor did not undergo proteolysis in the process (Figure 2c and 2d, respectively). In *Vibrio* 1F-97, production of TF_63_ or smORF_63_ led to rapid lysis of the bacterial culture and an accumulation of phage 63 particles in the cell-free culture fluids (Figure 2e and 2f, respectively, and Extended Data Figure 2a). These results reveal that phage 63 encodes a TF-smORF module near the *par* genes, and that smORF_63_, like smORF_72_, functions as an antirepressor.

**Figure 2.**
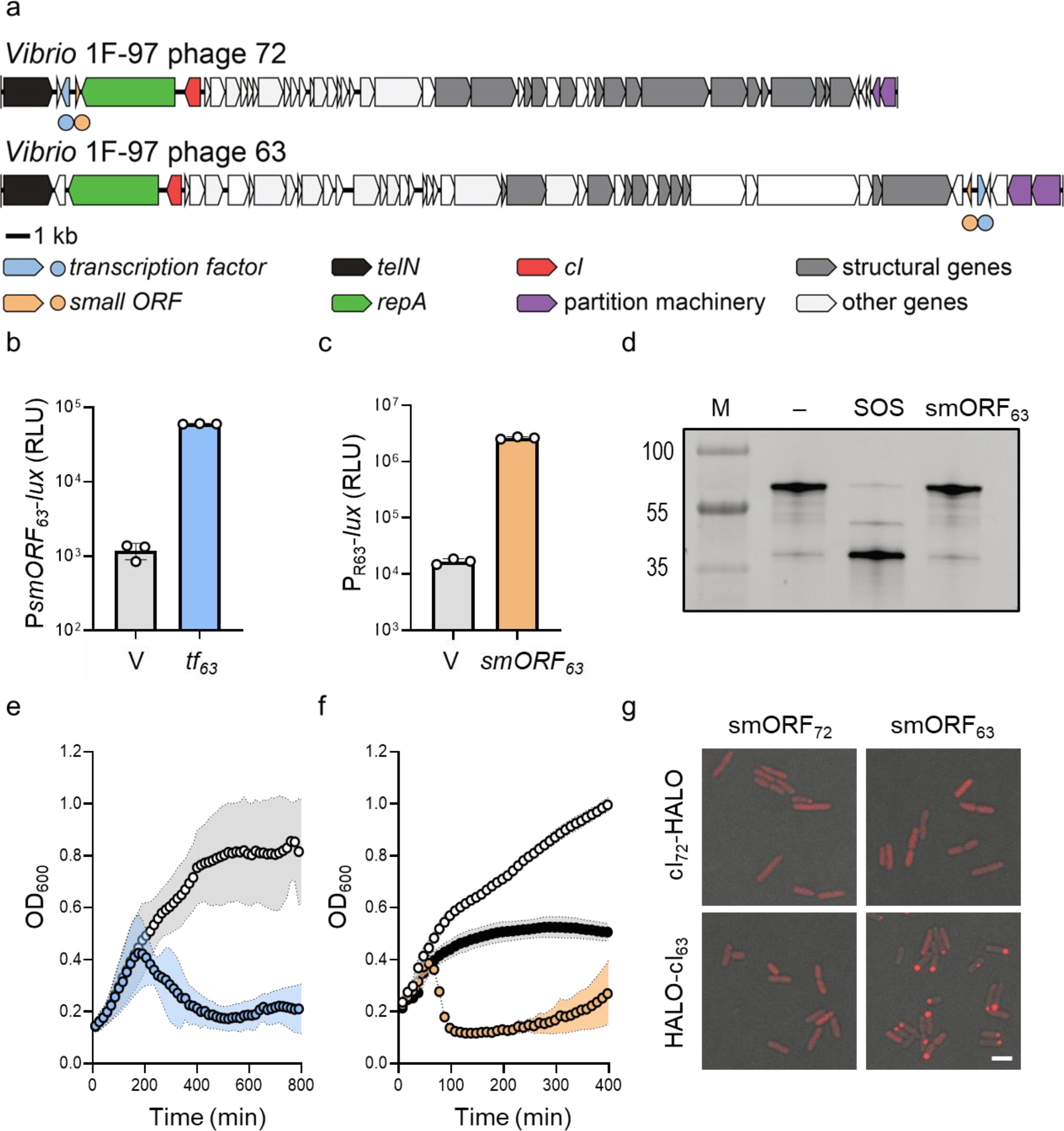
*Vibrio* 1F-97 is infected with two linear plasmid-like phages that control lysis via TF-smORF modules encoded in different genomic loci. **(a)** Organization of linear plasmid-like phage genomes in *Vibrio* 1F-97. Genes are colored by annotation as noted in the key below the figure. Circles below genomes denote the positions of the *tf* (blue) and *smORF* (orange) genes located at *repA-telN* (phage 72) or by *par* genes (phage 63). **(b)** P*smORF_63_*-*lux* expression from *E. coli* carrying an empty vector (V) or aTc-inducible *tf_63_* in medium containing aTc. **(c)** P_R63_-*lux* expression in *E. coli* carrying an empty vector (V) or aTc-inducible *smORF_63_* in medium containing aTc. The P_R63_-*lux* plasmid carries *cI_63_*, which natively represses reporter expression. **(d)** SDS-PAGE in-gel labeling of the phage 63 repressor (HALO-cI_63_) produced in *E. coli* carrying aTc-inducible *smORF_63_*. The treatments -, SOS, and smORF_63_ refer to water, ciprofloxacin, and aTc, respectively. M as in Figure 1b. **(e)** Growth of *Vibrio* 1F-97 carrying aTc-inducible *tf_63_* in medium lacking or containing aTc (white and blue, respectively). **(f)** Growth of *Vibrio* 1F-97 carrying aTc-inducible *smORF_63_* in medium containing aTc (orange), ciprofloxacin (black), or water (white). **(g)** Confocal microscopy of *E. coli* producing cognate and non-cognate pairs of smORF_72_ or smORF_63_ with cI_72_-HALO or HALO-cI_63_ labeled with HALO-TMR. Scale bar denotes 3 µm. All media contained aTc to induce *smORF* expression. Data are represented as means ± std with *n*=3 biological replicates (b, c, e, f). RLU as in Figure 1d (b, c). aTc; 50 ng mL^−1^ (b, c, d, e, f, g), ciprofloxacin; 500 ng mL^−1^ (d, f).

In phage VP882, the founding member of linear phages with TF-smORF modules, the Qtip antirepressor inactivates its partner cI repressor by sequestering it to the cell poles.^21^ To test whether this could be a general feature of these modules, we assessed the intracellular localization of the *Vibrio* 1F-97 phage repressor proteins. To do this, we engineered functional HALO-tag translational fusions to each cI (cI_72_-HALO and HALO-cI_63_). Figure 2g (top left) shows that cI_72_-HALO is largely diffuse in the cytoplasm when co-produced with smORF_72_. In contrast, HALO-cI_63_, which alone is diffuse in the cytoplasm, formed polarly localized puncta in the presence of smORF_63_ (Figure 2g, bottom right). Co-production of the two non-cognate pairs (smORF_72_ with HALO-cI_63_ and smORF_63_ with cI_72_-HALO) had no effect on localization of the repressors (Figure 2g bottom left and upper right, respectively), indicating that the smORF_63_-induced localization of HALO-cI_63_ is specific. Collectively, these results suggest that despite having equivalent pathway components, the two smORFs uniquely affect their target cI repressors.

### A search for phages with genes adjacent to partition genes reveals homoserine lactone-quorum-sensing TF-smORF modules

Our finding that phage genes encoding transcription factor-smORF modules are not restricted to *repA-telN* genomic regions motivated us to expand our search beyond this locus. Preliminary BLAST searches suggested that genes encoding homologs of *tf_63_* often occur near *parB* genes and within 25 Kb of conserved phage structural genes, mirroring what we show in Figure 2a. To systematically identify new TF-smORF modules, we searched NCBI for sequences encoding homologs of ParB and the major capsid protein from phage 63. We further filtered the output of that search for those hits that encode an apparent transcription factor within 10 Kb of the predicted *parB* gene. This search strategy revealed 56 unique contigs (Supplementary Table 1), approximately 90% of which (50/56) are putative linear plasmid-like phages on the basis of detectable *repA*-*telN* loci (Supplementary Table 3). We identified one *par*-associated node of interest that contained 5 phages (Figure 3a). Each phage carries an operon containing genes encoding an XRE-like DNA-binding protein and a LuxR-type transcription factor. LuxR-type proteins contain N-terminal acyl homoserine lactone (HSL) AI-binding domains and C-terminal helix-turn-helix DNA-binding domains.^22^ The *luxR* genes for two of the identified phages (ARM81ld of *Aeromonas* sp. ARM81 and Apop of *Aeromonas popoffii*) are associated with host strains that we had previously identified in a bioinformatic search for phage-encoded LuxR proteins.^23^ In that work, we showed that the phage LuxR proteins could be solubilized by the HSLs *Aeromonads* are known to produce, however, we were unable to deduce the functions of the phage LuxR proteins at the time due to an inability to obtain a bacterial isolate with the phage (*Aeromonas* sp. ARM81) and genetic intractability (*A. popoffi*). To expand our understanding of the roles of these phage modules, we successfully obtained *Aeromonas* sp. ARM81 and we developed genetic tools to study *A. popoffii*.

**Figure 3.**
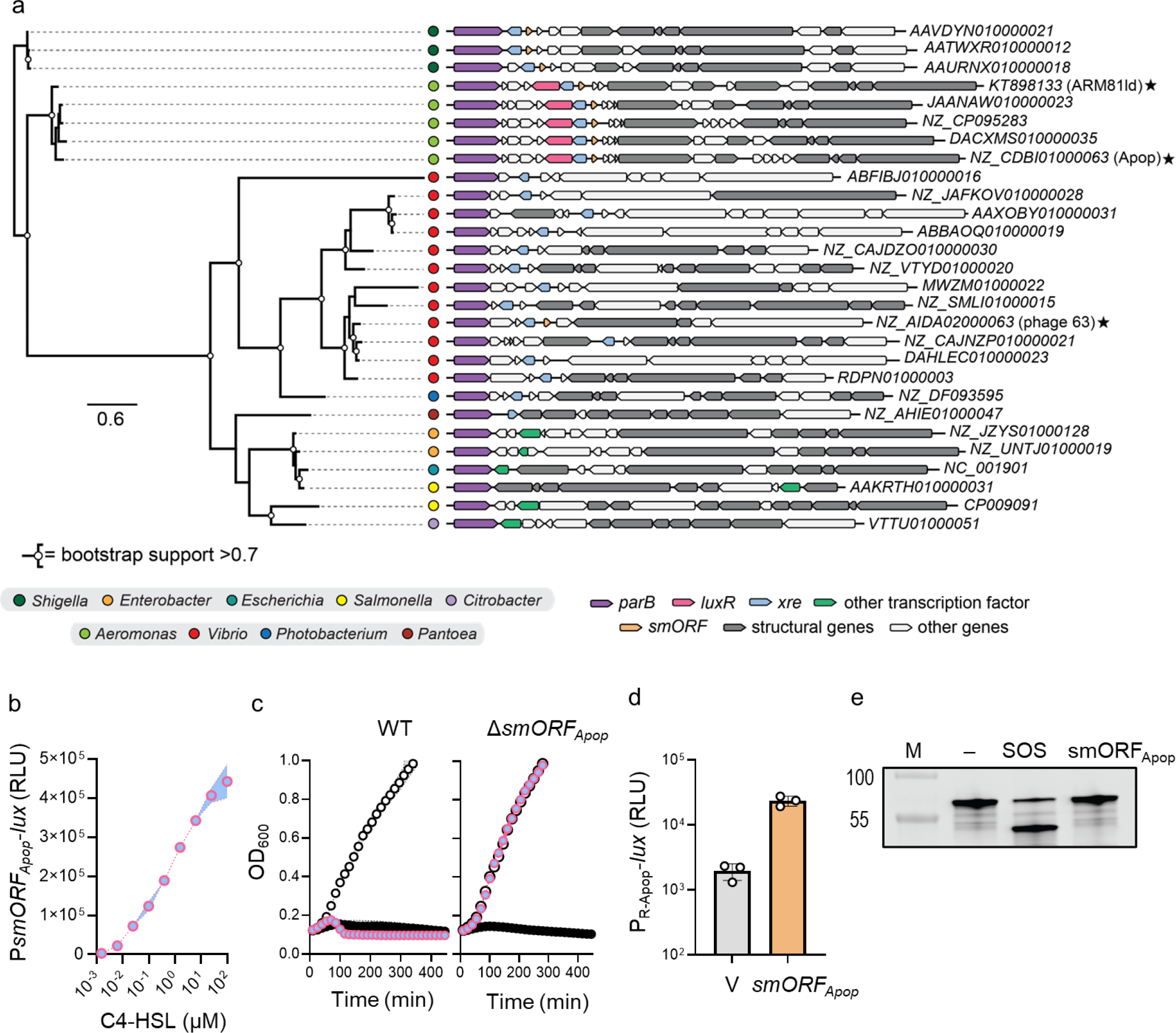
Phages with genes adjacent to *par* genes encode HSL-quorum-sensing-receptor TF-smORF modules that control lysis. **(a)** Phylogenetic tree (left) of 28 representative ParB proteins. ParB protein groups typically segregate with bacterial genera (colored circles, legend at bottom), but these groups are not always concordant with deeper taxonomic relationships. Nodes with bootstrap support >0.7 are indicated with white circles. Gene neighborhoods (right) for the 28 loci encoding *parB* (purple) and the major capsid protein (gray). ARM81ld, Apop, and phage 63 are denoted with stars. The *luxR*, *xre*, and *smORF* genes are colored pink, blue, and orange, respectively. Genes encoding predicted transcription factors that are not *luxR* or *xre* are colored in green. NCBI accession numbers are depicted to the right of each sequence. The scale bar indicates the number of amino acid substitutions per site. **(b)** P*smORF_Apop_*-*lux* expression from *E. coli* carrying aTc-inducible *xre_Apop_-luxR_Apop_* in medium with aTc and the indicated concentrations of C4-HSL. **(c)** Growth of *A. popoffii* carrying wild-type (WT) Apop (left) or ∆*smORF* Apop (right), each harboring aTc-inducible *xre_Apop_-luxR_Apop_* and grown in medium containing 5 ng mL^−1^ aTc (blue/pink), 1 μg mL^−1^ ciprofloxacin (black), or water (white). All media contained 10 µM C4-HSL. **(d)** P_R-Apop_-*lux* expression from *E. coli* carrying an empty vector (V) or aTc-inducible *smORF_Apop_* in medium containing aTc. The P_R-Apop_-*lux* plasmid carries two copies of *cI_Apop_* (see Methods) for native repression of reporter expression. **(e)** SDS-PAGE in-gel labeling of the Apop repressor (HALO-cI_Apop_) produced in *E. coli* carrying aTc-inducible *smORF_Apop_*. The treatments -, SOS, and smORF_Apop_ refer to water, ciprofloxacin, and aTc, respectively. M as in Figure 1b. Data are represented as means ± std with *n*=3 biological replicates (b, c, d). RLU as in Figure 1d (b, d). aTc; 50 ng mL^−1^ (b, d, e), ciprofloxacin; 500 ng mL^−1^ (e).

First, we investigated the *A. popoffii* XRE-LuxR (XRE_Apop_-LuxR_Apop_) pair. An unannotated 156 nt *smORF* resides approximately 150 nucleotides upstream of the *xre_Apop_-luxR_Apop_* operon near the *par* locus (Figure 3a). To determine if it is regulated by XRE_Apop_-LuxR_Apop_, we transformed a plasmid carrying aTc-inducible *xre_Apop_-luxR_Apop_* into *E. coli* harboring *lux* fused to the promoter of the candidate *smORF* (P*smORF_Apop_-lux*). Light production from P*smORF_Apop_-lux* commenced only when the *E. coli* reporter was supplied with C4-HSL, an AI natively produced by *Aeromonads* (Figure 3b). Thus, XRE_Apop_ and the LuxR_Apop_-AI complex control *smORF* expression. Consistent with this finding, deletion of the *smORF* locus from Apop abolished XRE_Apop_-LuxR_Apop_-AI-dependent lysis of *A. popoffii* but not ciprofloxacin-dependent lysis (Figure 3c). Analogous to what we show in Figures 1d and 2c, smORF_Apop_ is an antirepressor that prevents its partner repressor, cI_Apop_, from maintaining lysogeny (Figure 3d). Lastly, unlike DNA damage, smORF_Apop_ did not lead to cI_Apop_ autoproteolysis (Figure 3e), indicating that the XRE_Apop_-LuxR_Apop_-controlled module, as with those of the other phages studied so far, acts as an orthogonal SOS-independent pathway to induction.

To define the individual and combined contributions of XRE_Apop_ and LuxR_Apop_ to the activation of *smORF_Apop_* expression, we generated expression vectors carrying only pTet-*xre_Apop_* or only pTet-*luxR_Apop_* and tested them in the P*smORF_Apop_-lux* assay. Neither XRE_Apop_ nor LuxR_Apop_ alone drove reporter output irrespective of the presence of C4-HSL, indicating that in addition to AI, *smORF_Apop_* expression depends on both transcription factors (Figure 4a). To validate these findings, we engineered *xre_Apop_*-P*smORF_Apop_*-*lux* and *luxR_Apop_*-P*smORF_Apop_*-*lux* reporter plasmids, which carry the identical P*smORF_Apop_-lux* reporter sequence but that include either full-length *xre_Apop_* or full-length *luxR_Apop_*, controlled by the native promoter. We introduced pTet*-xre_Apop_* or pTet*-luxR_Apop_* into *E. coli* harboring the reporters. Heterologous *xre_Apop_* expression induced light production only when *luxR_Apop_* was present on the reporter plasmid and AI was supplied (Figure 4b and 4c). By contrast, heterologous *luxR_Apop_* expression did not induce light production in either case, irrespective of the presence of AI (Figure 4b and 4c). We interpret these results as follows: XRE_Apop_ activates expression of the *xre_Apop_-luxR_Apop_* operon, driving both XRE_Apop_ and LuxR_Apop_ production. Together, LuxR_Apop_, bound to C4-HSL, and XRE_Apop_, are required to activate P*smORF_Apop_* expression, launching the phage lytic cascade and host-cell lysis (Figure 4d). Confirming the above regulatory arrangement *in vivo*, RT-qPCR revealed that overexpression of *xre_Apop_* in *A. popoffii* led to rapid activation of expression of *xre_Apop_* and *luxR_Apop_*, whereas overexpression of *luxR_Apop_* did not (Figure 4e).

**Figure 4.**
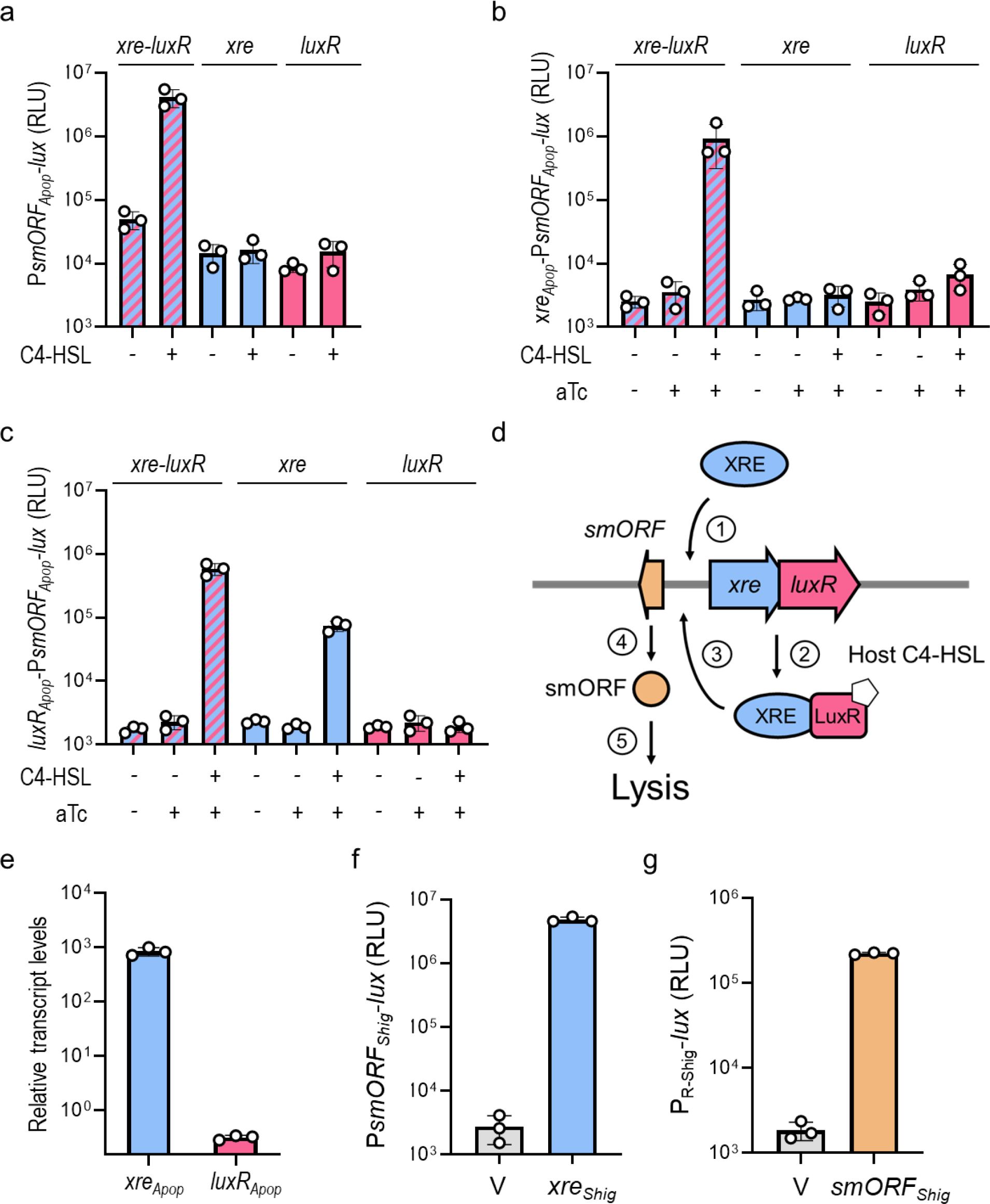
Activation of P*smORF_Apop_* requires XRE_Apop_, LuxR_Apop_, and the C4-HSL AI. **(a)** P*smORF_Apop_*-*lux* activity from a plasmid in *E. coli* carrying a second vector with either aTc-inducible *xre_Apop_-luxR_Apop_*, *xre_Apop_* alone, or *luxR_Apop_* alone in medium lacking or containing C4-HSL (− and +, respectively). All media contained aTc. **(b)** Expression of P*smORF_Apop_*-*lux* from a plasmid carrying *xre_Apop_* under its native promoter in *E. coli* carrying a second plasmid encoding aTc-inducible *xre_Apop_-luxR_Apop_*, *xre_Apop_* alone, or *luxR_Apop_* alone. Media contained the indicated combinations of water (−) or aTc (+), and DMSO (−) or C4-HSL (+), as indicated below the axis. **(c)** Expression of P*smORF_Apop_*-*lux* from a plasmid carrying *luxR_Apop_* under its native promoter in *E. coli* carrying aTc-inducible *xre_Apop_-luxR_Apop_*, *xre_Apop_* alone, or *luxR_Apop_* alone. Treatments as in (b). **(d)** Proposed model for regulation of the TF-smORF module in Apop. (1) XRE_Apop_ activates expression of the *xre_Apop_-luxR_Apop_* operon, and (2) increased production of LuxR_Apop_ and XRE_Apop_ occurs. (3) LuxR_Apop_ when bound to the C4-HSL AI ligand (pentagon), together with XRE_Apop_, activates expression of the counter-oriented *smORF_Apop_* gene, and also activates expression of *xre_Apop_*-*luxR_Apop_*. (4) smORF_Apop_ inhibits the cI_Apop_ repressor, (5) leading to host-cell lysis. The co-requirement for AI-bound LuxR_Apop_ and XRE_Apop_ could be due to a protein-protein interaction or an alteration in the topology of the XRE_Apop_ DNA-binding site in favor of expression of *smORF_Apop_*. The mechanism remains to be determined. **(e)** Relative transcript levels of the *xre_Apop_-luxR_Apop_* operon from the Apop genome following plasmid expression of either aTc-inducible *xre_Apop_* or *luxR_Apop_* in *A. popoffii*. All media contained C4-HSL and aTc. Primer pairs specific to the intergenic region in the *xre_Apop_-luxR_Apop_* locus but absent from the aTc-inducible *xre_Apop_* and *luxR_Apop_* plasmids were used to measure native *xre_Apop_-luxR_Apop_* expression (see Methods). Relative transcript levels are the amount of *xre_Apop_-luxR_Apop_* DNA relative to the amount of *rpoB* DNA, normalized to the sample overexpressing *luxR_Apop_*. **(f)** P*smORF_Shig_*-*lux* expression in *E. coli* carrying an empty vector (V) or aTc-inducible *xre_Shig_* in medium containing aTc. **(g)** P_R-Shig_-*lux* expression in *E. coli* carrying an empty vector (V) or aTc-inducible *smORF_Shig_* in medium containing aTc. The P_R-Shig_-*lux* plasmid carries *cI_Shig_*, which natively represses reporter expression. Data are represented as means ± std with *n*=3 biological replicates (a, b, c, f, g) and as means ± std with *n*=3 biological replicates and *n*=4 technical replicates (e). RLU as in Figure 1d (a, b, c, f, g). aTc; 50 ng mL^−1^ (a, b, c, f, g), aTc; 5 ng mL^−1^ (e), C4-HSL; 10 µM (a, b, c, e).

### Variations in host bacterial signal transduction cascades may affect which transcription factor is employed by the phage

We wondered whether the XRE and LuxR components always co-occur in phage TF-smORF modules. A tBLASTn search using XRE_Apop_ as the query revealed three contigs, likely from linear plasmid-like phages, in different *Shigella sonneii* genomes (Figure 3a and Extended Data Figure 3a). As in the *Aeromonas* phages, the *Shigella* phage *xre (xre_Shig_*) genes are located downstream of the *par* genes and oriented divergently to a hypothetical 165 nt *smORF*. Unlike in the *Aeromonas* phages, these putative *Shigella* phages lack *luxR* genes. *Shigella* are not known to produce HSL AIs, thus a LuxR component might not provide benefits to the phage in the *Shigella* host.

We could not obtain the *S. sonneii* isolates to determine if the phage XRE_Shig_ and smORF_Shig_ components are functional *in vivo*. To circumvent this issue, we synthesized a P*smORF_Shig_-lux* reporter and an inducible *xre_Shig_* gene. Production of XRE_Shig_ activated P*smORF_Shig_-lux* in recombinant *E. coli*, demonstrating that this module does not require a partner LuxR (Figure 4f). We also synthesized inducible *smORF_Shig_* and the predicted cI_Shig_-repressed promoter (*cI_Shig_*-P_R-Shig_-*lux*) to assess whether our predicted smORF_Shig_ is an antirepressor. Indeed, expression of *smORF_Shig_* in recombinant *E. coli* derepressed *cI_Shig_*-P_R-Shig_-*lux* (Figure 4g). Taken together, these results reveal that *Shigella* phages encode TF (i.e., XRE)-smORF lysis modules. More generally, these results suggest that the particular sets of regulatory components encoded on different plasmid-like phages have been shaped through evolution, presumably by relevant host-sensory cues.

### Phage-encoded transcription factor-small protein modules dictate which prophage is induced in polylysogenic hosts

The literature on polylysogeny predicts that inter-prophage competition arises when co-occurring prophages are induced by the same trigger (e.g., DNA damage).^24^ Our above discovery and characterization of additional phage lysis pathways suggest that the outcomes of such competitions could depend on the inducer responsible for launching lysis. To investigate this possibility, we used *Vibrio* 1F-97 as a model given that it is naturally polylysogenic for two linear plasmid-like prophages, each possessing its own additional sensory pathway. Regarding similarity between the pathways, we note that TF_63_ shares ~53% amino acid identity to TF_72_ while their partner smORFs share only 9 identical residues in pairwise alignment (11% amino acid identity; Figure 2a). We assessed whether either phage TF could cross-activate expression of its non-cognate *smORF* gene. Figure 5a shows that TF_72_ did not activate light production from a P*smORF_63_-lux* reporter, and similarly, TF_63_ did not activate light production from a P*smORF_72_-lux* reporter. Consistent with our earlier finding that the smORFs did not affect localization of their non-cognate repressors (Figure 2g), neither non-cognate pair (pTet-*smORF_63_* with *cI_72_*-P_R72_-*lux*; pTet-*smORF_72_* with *cI_63_*-P_R63_-*lux*) generated P_R_-driven light production (Figure 5b). Thus, each phage-encoded module is specific at the level of *smORF* expression and smORF-driven repressor inactivation.

**Figure 5.**
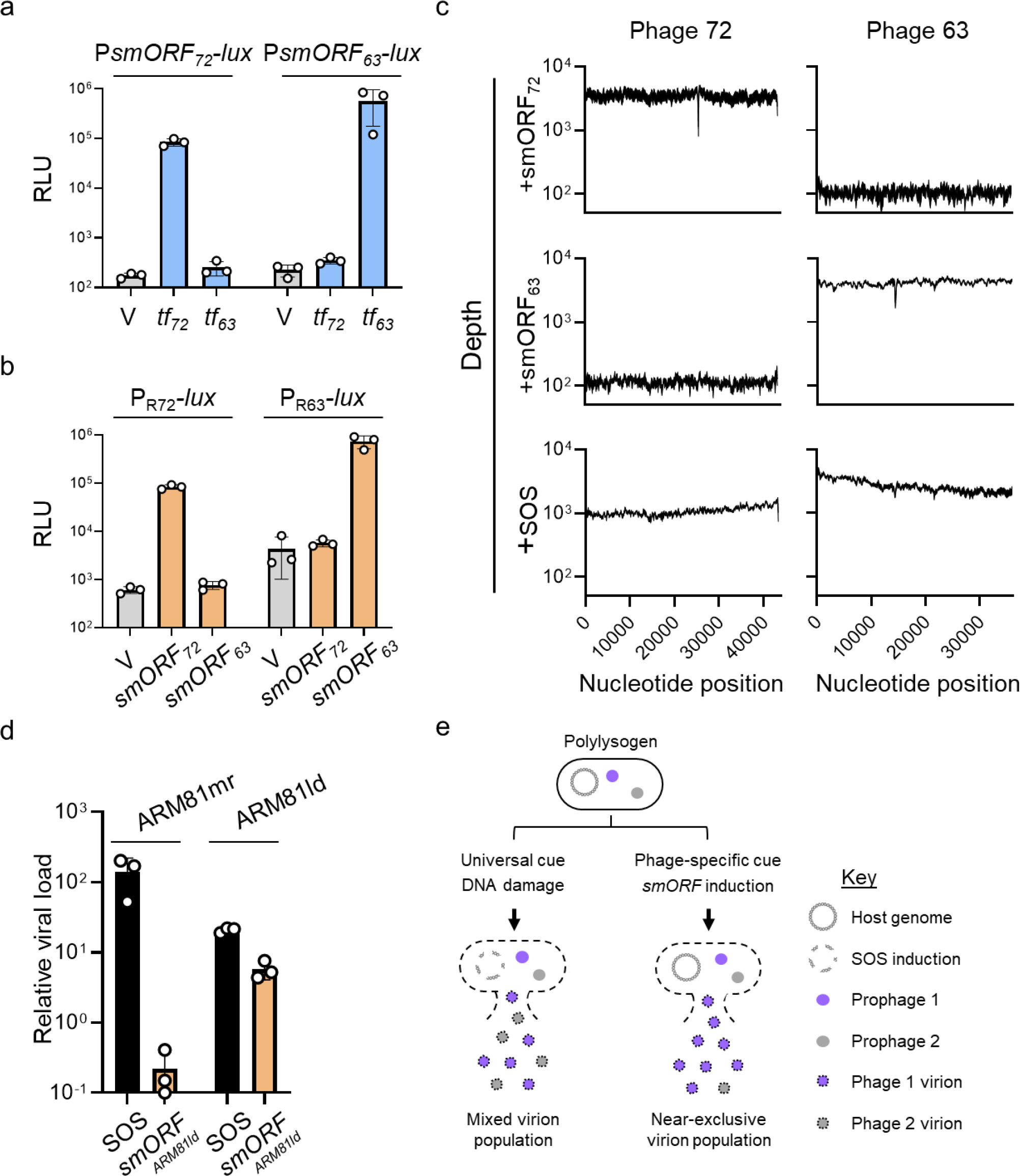
Phage-encoded TF-smORF modules exclusively drive self-induction in polylysogenic hosts. **(a)** P*smORF*-*lux* output from *E. coli* harboring P*smORF_72_*-*lux* or P*smORF_63_*-*lux* and a second plasmid encoding either an empty vector (V), aTc-inducible *tf_72_* or aTc-inducible *tf_63_*, each grown in medium containing aTc. **(b)** Light production from *E. coli* harboring cI-repressed P_R72_-*lux* or P_R63_-*lux* reporter plasmids and a second plasmid encoding either an empty vector (V), aTc-inducible *smORF_72_* or aTc-inducible *smORF_63_*, each grown in medium containing aTc. **(c)** Whole-genome sequencing of viral particles prepared from *Vibrio* 1F-97 cultures carrying aTc-inducible *smORF_72_* or aTc-inducible *smORF_63_* grown in medium containing aTc or ciprofloxacin for smORF induction or activation of the SOS response, respectively. Depth refers to the number of read counts. **(d)** Detection of viral particles prepared from *Aeromonas* sp. ARM81 cultures carrying aTc-inducible *smORF_ARM81ld_* grown in medium containing ciprofloxacin for SOS or aTc for smORF_ARM81ld_ production. All media contained C4-HSL. Relative viral load is the amount of ARM81mr- or ARM81ld-specific DNA (*cI_ARM81mr_* and *cI_ARM81ld_*, respectively) in the induced samples relative to an uninduced sample measured by qPCR. **(e)** Proposed model for how TF-smORF modules influence inter-prophage competition outcomes in polylysogenic bacteria. Left side: Exposure of a polylysogen to a ubiquitous trigger (i.e., SOS activation/DNA damage) fosters competitive conditions in which multiple prophages share host cell resources to replicate and produce viral particles. Right side: Prophage-specific induction via a TF-smORF module (the purple inducer is present in this example) leads to non-competitive conditions in which only the induced phage garners host cell resources leading to almost exclusive production of that phage particle. Data are represented as means ± std with *n=3* biological replicates (a, b) and as means ± std with *n*=3 biological replicates and *n*=4 technical replicates (d). RLU as in Figure 1d (a, b). aTc; 50 ng mL^−1^ (a, b, c), aTc; 5 ng mL^−1^ (d), ciprofloxacin; 500 ng mL^−1^ (c), ciprofloxacin; 1 µg mL^−1^(d), C4-HSL; 10 µM (d).

*Vibrio* 1F-97 cultures lyse when induced with ciprofloxacin, smORF_63_, or smORF_72_ (Figures 1c and 2f). Our results defining pathway specificity predict that the population of phage particles produced could vary depending on which pathway drives lysis. To test this hypothesis, we performed whole genome sequencing of phage particle preparations in *Vibrio* 1F-97 cultures induced by ciprofloxacin, smORF_63_, or smORF_72_. Figure 5c shows that ciprofloxacin treatment drove production of DNA corresponding to both phage 72 and phage 63, whereas cultures induced by smORF_72_ or smORF_63_ resulted in near-exclusive production of phage 72 particles and phage 63 particles, respectively. Thus, the induction cue dictates the distribution of phage particles produced by the polylysogenic host.

To test the generality of the above finding, we turned to *Aeromonas* sp. ARM81 harboring the ARM81ld prophage with the *par*-associated *xre_ARM81ld_-luxR_ARM81ld_*-*smORF_ARM81ld_* module. *Aeromonas* sp. ARM81 is polylysogenic and contains an integrated prophage, ARM81mr, in addition to the plasmid-like ARM81ld prophage.^25^ We have no genomic evidence for an additional sensory pathway on ARM81mr phage. Similar to Apop, the ARM81ld module depends on the C4-HSL AI to launch lysis, and its partner small ORF (encoded by the 162 nt *smORF_ARM81ld_* gene) does not proteolyze its partner cI repressor, unlike what occurs following DNA damage (Extended Data Figure 3b-e). Previous work indicates that both ARM81ld and ARM81mr are induced by DNA damage but that most particles produced are of ARM81mr.^25^ Consistent with the earlier report, when ciprofloxacin was administered, there was a 140-fold increase in ARM81mr phage particles and only a 22-fold increase in ARM81ld particles, relative to when no inducer was added (Figure 5d). In contrast, induction of expression of *xre_ARM81ld_-luxR_ARM81ld_* with C4-HSL led to a 5-fold increase in ARM81ld particles, whereas ARM81mr particles were reduced 5-fold compared to their the uninduced levels (amounting to a 700-fold reduction in ARM81mr particles compared to ciprofloxacin treatment; Figure 5d). Based on our results with both *Vibrio* 1F-97 and *Aeromonas* sp. ARM81, we conclude that while DNA damage results in induction and competition between co-residing prophages, possession of an additional sensory pathway can drive host-cell lysis exclusively by the phage that encodes the module (Figure 5e).

## DISCUSSION

Despite most bacteria being predicted to be polylysogenic, an understanding of the effects of polylysogeny both on the hosts and on their resident phages has been constrained by the limited models available and a lack of known prophage induction cues beyond the SOS response. Consequently, competition among co-residing temperate prophages is generally considered a ‘sprint’ in which, in response to a single trigger (i.e., the SOS response), differences in particular phage properties (e.g., replication rates, packaging rates, burst size, etc.), dictate how many particles of each phage are produced. Our present work shows that in polylysogens, prophage induction via SOS-independent pathways generates conditions in which differential phage induction can occur and the outcome varies depending on the induction cue. While tuning into the host SOS response remains a universal mechanism by which prophages perceive host-cell stress, the additional sensory pathways we discover suggest that more specialized conditions exist which favor induction of one phage over another.

Our results show that phage 72 and phage 63 of *Vibrio* 1F-97 do not cross-activate each other’s lytic programs. This specificity is maintained at the level of *smORF* expression and smORF-driven repressor inactivation. By contrast, investigation of polylysogenic *Salmonella* strains revealed that some phage antirepressor proteins can inactivate cognate and non-cognate repressors, thus enabling synchronization of induction of multiple prophages.^26^ This latter arrangement is thought to be vital for prophages with slow induction responses, allowing them to ‘piggyback’ off of prophages that are induced more rapidly.^26^ The finding that prophage induction by an additional cue does not trigger induction of co-residing prophages could be a strategy that proves especially successful when host resources are limiting because it ensures exclusive reproduction and dissemination of only the induced phage.

The genes encoding the phage transcription factors TF_72_, TF_63_, XRE_Apop_-LuxR_Apop_, and XRE_ARM81ld_-LuxR_ARM81ld_ all required synthetic induction to drive lysis, a situation that exactly parallels our findings for VqmA_Phage_ in phage VP882.^9^ We compare this scenario to DNA-damaging compounds that are universally used to induce prophages in the laboratory. DNA-damaging compounds do not generally occur in nature at the concentrations used in experiments. Thus, we argue that the identities of the natural cues that drive lysis even in intensively-studied (i.e., SOS-dependent) prophages remain mysterious. Presumably, the native cues will reflect the habitats in which the bacterial hosts reside. For example, some cyanophages, by encoding photosystem components, increase the light harvesting capacities of their hosts.^27^ Other phages associated with animals have capsids decorated with Ig-like domains that enable phage particles to adhere tightly to mucus, creating a reservoir of virions that can infect incoming naïve bacteria.^28^ In so doing, these phages confer protection to the host animal from bacterial infection. These few examples, strongly suggest that the signals that induce the additional pathways to lysis revealed here, as well as others that are identified going forward, will be particular to each host-phage partnership and the niche in which they reside. We propose that, while discovered long after the SOS cue, some or all of these pathways may in fact be the primary determinants of phage lifestyle transitions and the arbiters of inter-prophage competition in real-world settings.

