## Supplementary material for "Small protein modules dictate prophage fates during polylysogeny": si

### METHODS

**Sequence Retrieval, RepA/TelN and ParB-associated loci**

To identify examples of phage genomes with convergently-oriented *telN* and *repA* genes, we examined sequences from the following databases: NCBI nt, IMG/VR v3 (specifically, the file 'IMGVR_all_nucleotides.fna'),[1] Cenote Human Virome Database v1.1 (CHVD_clustered_mash99_v1.fna),[2] Global Ocean Virome 2 database (GOV2_viral_populations_larger_than_5Kb_or_circular.fna),[3] the Gut Phage Database (GPD_sequences.fna),[4] a curated set of linear plasmid-phages (retrieved via NCBI accession numbers in Table S4 of Pfeifer *et al*),[5] and the Metagenomic Gut Virus Catalog (mgv_contigs.fna).[6] In February of 2022, most databases were searched using both tBLASTn and profile-HMM-based search strategies (the exception being NCBI nt, which was too large to annotate anew using profile HMMs). A manual examination of established lysis-control loci[7] revealed that the associated *telN* gene always encoded a protein that matched closely to Pfam profile PF16684, whereas the following set of seven families encompassed the diversity of observed *repA* sequences: PF13362, PF02399, PF08707, PF10661, PF02502, and PF13604. Thus, we extracted these seven families from Pfam (v34)[8] into a custom HMM database. We then used the gene finder MetaGeneMark[9] to predict open reading frames (ORFs) from the aforementioned nucleotide files using default parameters. The resulting proteins were used in a profile HMM search with HMMER3[9] against the *repA*/*telN* custom database and nucleotide sequences were extracted that contained both *telN* and *repA* genes, only if they were convergently oriented and within 10 Kb of one another. To complement this search strategy, we also used the predicted TelN and RepA proteins from vibriophage VP882 in a tBLASTn search against these nucleotide databases (accession numbers YP_001039865 and YP_001039868, respectively). We retained only the hits with e-values better than 0.001 which also covered > 50% of the query sequence. For consistency, we re-annotated all retrieved sequences using a common method, as follows. We used the gene-finder MetaGeneMark[10] to predict open reading frames (ORFs) using default parameters. We next used their amino acid sequences in a profile HMM search with HMMER3[9] against TIGRFAM[11] and Pfam[8] profile HMM databases. The highest scoring profile was used to annotate each ORF. As above, we further refined our database by considering only contigs with convergently oriented *telN* and *repA* genes within 10 Kb of one another. The HMM and tBLASTn-based search strategies produced highly redundant (but not identical) sequence sets, as the same databases were used and many identical sequences are listed across multiple databases. Thus, we dereplicated our combined sequence files using cd-hit-est with the following parameters: '-c 1.0 -aS 1.0 -g 1 -d 0'. We manually examined the DNA sequence located between each *telN* and *repA* gene from this dereplicated set, extracting the intervening nucleotides to a second locus-specific dataset. We dereplicated these sequences as described above to produce the final set of 274 loci referenced in the text and detailed in Supplementary Table 1. The NCBI nucleotide accession that encodes phage 63 (NZ_AIDA02000063) is dated 20-March-2022, after our initial February search. The sequence for phage 72 appears more than once in the NCBI nucleotide database (NZ_AIDA02000072, dated 20-March-2022, and KP795532, dated 18-Nov-2019). Presumably, these similar entries result from separate sequencing analyses of identical (or similar) bacteria. For these reasons, our February BLAST search only initially revealed phage 72, while phage 63 was discovered subsequently.

To identify examples of putative lysis-control loci associated with the *parB* gene, we first performed a BLASTp search against the NCBI nr/nt database using the ParB protein from phage 63 as a query (accession number WP_016786072). From the top 10,000 proteins revealed by this search, we retrieved the corresponding nucleotide file in NCBI and examined the locus surrounding *parB* by downloading DNA +/- 50 Kb from the gene boundaries (this strategy retrieved the full contig sequence for many of the associated nucleotide files). To identify *parB* genes associated with the phage structural locus present in phage 63, we used tBLASTn to query these sequences with the major capsid protein from this phage (accession number WP_016786053), which is one of the most conserved phage protein folds.[12] We filtered for hits with over 25% amino acid identity across 70% of the query sequence, revealing 121 putative *parB* loci present on 118 unique sequences. These 121 loci were dereplicated as described for *telN*/*repA* and filtered for the presence of a full-length *parB* gene and a predicted transcription factor within 10 Kb of *parB*. This analysis produced 56 sequences, which are referenced in Supplementary Table 1.

**Phylogenetic analysis of TelN and ParB**

From a manual examination of the retrieved RepA- and TelN-encoding sequences, we observed that all TelN proteins shared the same profile HMM as their top-scoring annotation (PF16684.8). This pattern stood in contrast to predicted RepA-encoding sequences, which hit best to one of six different protein families. The consistent TelN annotation suggested to us that this protein may be conserved despite extensive sequence divergence between the phage genomes considered, and thus, it provided a good phylogenetic bellwether for this locus. Indeed, the set of non-redundant, full-length TelN proteins from these datasets aligned well using MUSCLE with default parameters.[13] A preliminary phylogenetic tree was generated from these sequences using the Geneious Tree Builder per the UPGMA clustering method and with a Jukes-Cantor distance model. From this tree, a subset of 34 sequences was selected to best represent the full TelN phylogenetic tree and the genetic diversity encoded by its associated gene neighborhoods. Figure 1A depicts a phylogenetic tree generated from these 34 representative sequences, which was produced using PhyML with the LG substitution model and bootstrapped 100 times. A similar procedure was used to produce the ParB phylogeny depicted in Figure 3A. In this case, a preliminary phylogenetic tree was produced from the set of 56 ParB proteins in the full dataset using the Geneious Tree Builder per the UPGMA clustering method and with a Jukes-Cantor distance model. From this tree, a subset of 28 sequences was selected to best represent the full

ParB phylogenetic tree and the genetic diversity encoded by its associated gene neighborhoods. This final ParB tree was produced using PhyML with parameters identical to those used for TelN.

##### Prediction of autoproteolytic cI proteins

All 271 predicted cI proteins were used to perform a batch search of NCBI's Conserved Domain (CD) database. The output of this search included conserved domains and other protein features for each cI protein queried. Among these features were predicted catalytic sites that corresponded to known catalytic residues in canonical autoproteolytic cI proteins.[14] Thus, we used this feature table to define the 61 predicted autoproteolytic cI proteins and labeled the remaining 210 proteins as putatively non-autoproteolytic. The clustering analysis in Supplementary Table 2 was performed using the DNA sequences between *repA* and *telN* genes and grouped sequences that shared 80% nucleotide identity over 95% the length of the shorter sequence in the same cluster. We used the program 'cd-hit-est' with the following parameters: -c 0.8 -aS 0.95 -g 1 -d 0.

##### Phylogenetic analysis of linear plasmid-like phages encoding partition-associated *xre* modules in *Shigella* and *Aeromonas*

NCBI BLASTp of the $XRE_{Apop}$ sequence used as a representative of the three *Aeromonas* phages identified in the *par*-associated analysis (detailed above) retrieved three predicted linear plasmid-like phages in *Shigella*. For each phage genome, genes were called using Prodigal 2.6.3,[15] and gene diagrams were constructed using custom python scripts. Genes were annotated using Prokka 1.11,[16] and annotations were supplemented with NCBI BLASTp searches by hand.[17] Phage genome tree (Extended Data Figure 3a) was constructed using VICTOR,[18] yielding an average support of 78%. The numbers above branches are pseudo-bootstrap support values from 100 replications.

##### Bacterial strains and growth conditions

*E. coli* and *Aeromonads* were grown with aeration in Luria-Bertani (LB-Miller, BD-Difco) broth. *Vibrio* strains were grown in LB with 3% NaCl. All strains were grown at 30° C. Strains used in the study are listed in Supplementary Table 4. Unless otherwise noted, antibiotics, were used at: 100 μg $mL^{-1}$ ampicillin (Amp, Sigma), 50 μg $mL^{-1}$ kanamycin (Kan, GoldBio), and 5 μg $mL^{-1}$ chloramphenicol (Cm, Sigma). Inducers were used as follows: *E. coli*: 200 μM isopropyl beta-D-1-thiogalactopyranoside (IPTG, GoldBio), and 50 ng $mL^{-1}$ anhydrotetracycline (aTc, Clontech). *Vibrios*: 500 ng $mL^{-1}$ ciprofloxacin (Sigma) and 50 ng $mL^{-1}$ aTc. *Aeromonads*: 1 μg $mL^{-1}$ ciprofloxacin and 5 ng $mL^{-1}$ aTc. C4-HSL was supplied at a final concentration of 10 μM except for the experiment shown in Figure 3b, in which it was administered at the indicated concentrations.

**Cloning techniques**

All primers and dsDNA (gene blocks) used for plasmid construction and qPCR, listed in Supplementary Table 5, were obtained from Integrated DNA Technologies or Twist Bioscience. Gibson assembly, intramolecular reclosure, and traditional cloning methods were employed for all cloning. PCR with iProof was used to generate insert and backbone DNA. Gibson assembly relied on the HiFi DNA assembly mix (NEB). The Apop- and ARM81ld-based $P_R$-*lux* reporter constructs in *E. coli* (JSS-3346k and JSS-3348k, respectively) required the addition of a second copy of the *cI* repressor under its native promoter in the plasmid for function (see Table 5). All enzymes used in cloning were obtained from NEB. Plasmids used in this study are listed in Supplementary Table 6. Transfer of plasmids into *Vibrio* 1F-97, *A. popoffii*, and *Aeromonas* sp. ARM81 was carried out by conjugation followed by selective plating on TCBS agar supplemented with Kan, LB plates supplemented with Amp and Kan, and LB plates supplemented with Kan and Cm, respectively.

**Growth, lysis, and reporter assays**

Overnight cultures were back-diluted 1:100 with fresh medium with appropriate antibiotics prior to being dispensed (200 µL) into 96 well plates (Corning Costar 3904). Cells were grown in the plates for 90 min before ciprofloxacin, aTc, or C4-HSL were added as specified. Wells that did not receive treatment received an equal volume of water or DMSO. A BioTek Synergy Neo2 Multi-Mode reader was used to measure $OD_{600}$, or $OD_{600}$ and bioluminescence. Relative light units (RLU) were calculated by dividing the bioluminescence readings by the $OD_{600}$ at that time.

**RT-qPCR**

Overnight cultures were back-diluted 1:100 and grown for 90 min before administration of aTc at the indicated concentrations. In Figure 4e, cells were collected 15 min after aTc induction. In Extended Data Figure 1e, cells were collected at 0 min and 90 min after aTc induction. Harvested cells were treated with RNAProtect Bacteria Reagent (Qiagen) according to the supplier's protocol. Total RNA was isolated from cultures using the RNeasy Mini Kit (Qiagen). RNA samples were treated with DNase using the TURBO DNA-free Kit (Thermo). cDNA was prepared as described using SuperScriptIII Reverse Transcriptase (Thermo). SYBR Green mix (Quanta) and Applied Biosystems QuantStudio 6 Flex Real-Time PCR detection system (Thermo) were used for real-time PCR. Each cDNA sample was amplified in technical quadruplicate and data were analyzed by a comparative CT method which the indicated target gene was normalized to an internal bacterial control gene (*rpoB*).

**qPCR and viral preparation**

Viral preparations consisted of non-chromosomal DNA (RQ1, RNase-Free DNase, Promega) prepared from 1 mL of cells of the indicated strains. Overnight cultures were back-diluted 1:100 and grown for 90 min before being divided into 3 equal volumes and exposed to treatments as specified. Cultures were grown for an additional 5 h prior to collection of cell-free culture fluids (Corning SpinX). qPCR reactions were performed as described above for RT-qPCR reactions. 1

μL of purified non-chromosomal DNA was used for each qPCR reaction. Data were analyzed by normalizing the CT values of samples treated with ciprofloxacin or aTc to the CT values of samples treated with water using a primer set to the indicated phage. Viral preparations for whole genome sequencing (SeqCenter) were prepared exactly as described for qPCR, except by column purification (Phage DNA Isolation Kit, Norgen Biotek).

**Confocal microscopy**

Confocal microscopy of HALO-tagged cI proteins with untagged smORFs was carried out as described,[19] with minor modifications. Briefly, overnight cultures of *E. coli* T7Express lysY/$I^q$ carrying the indicated HALO fusion and inducible *smORF* vector were diluted 1:200 in medium containing 50 ng $mL^{-1}$ aTc and 1 μM HALO-TMR. Cultures were grown for an additional 3 h, subjected to centrifugation (16,100 x g for 1 min), washed twice with PBS, and resuspended in fresh PBS to a final $OD_{600}$ = 0.1-0.3. 5 μL of each sample was spotted onto a glass coverslip and overlaid with an LB agar pad. Samples were imaged approximately 45 min later using a Leica SP8 Confocal microscope. HALO-TMR was excited with 561 nm light and detected within the range 569-625 nm.

***in vitro* HALO-cI repressor cleavage and in-gel HALO detection**

Assessment of cleavage of HALO-cI proteins in response to DNA damage or smORF induction was carried out in *E. coli* according to a previously described method,[7] with minor modifications. Briefly, overnight cultures of *E. coli* T7Express lysY/$I^q$ carrying the indicated HALO fusion plasmid and cognate, aTc-inducible smORF vector were diluted 1:200 in medium and grown for 2.5 h with shaking. 200 μM IPTG was added to the cultures before being divided into 3 equal volumes followed by administration of the relevant treatment as specified. The treated cultures were incubated without shaking for an additional 2.5 h. Cells were collected by centrifugation (16,100 x g for 1 min), resuspended in BugBuster containing 1 μM HALO-$Alexa_{660}$ (excitation/emission:

663/690 nm). The cleared supernatant, collected after centrifugation of the lysate (16,100 x g for 10 min), was loaded onto a 4-20% SDS-PAGE stain-free gel. Gels were imaged using an ImageQuant LAS 4000 imager under the Cy5 setting for HALO-Alexa$_{660}$ before being exposed to UV-light for 7 min and re-imaged under the EtBr setting for total protein. Exposure times never exceeded 30 sec.

**Quantitation and statistical analyses**

Software used to collect and analyze data generated in this study consisted of: GraphPad Prism 9 for analysis of growth and reporter-based experiments; Gen5 for collection of growth and reporter-based data; Geneious Prime 2020 and SnapGene v6 for analysis of publicly available data and primer design; QuantStudio for qPCR collection; LASX for acquisition of confocal micrographs; and FIJI for image analyses. Data are presented as the means ± std. The number of technical and independent biological replicates for each experiment are indicated in the figure legends.

**Data and software availability**

Growth and reporter data presented in each panel of this study is available in Supplementary Table 7. Unprocessed gels and micrographs from this study are deposited on Zenodo (doi: 10.5281/zenodo.7083051). Other experimental data that support the findings of this study will be provided without restriction by request from the corresponding author.

### METHODS REFERENCES

#### ACKNOWLEDGMENTS

We thank Professor Matin Polz (University of Vienna) for gifting us 1F-97 and for thoughtful discussions throughout. We thank Julie Chen (Broad Institute), Abhishek Biswas (Princeton University), and all members of the Bassler lab for insightful discussions. This work was supported by the Howard Hughes Medical Institute, National Science Foundation grant MCB-1713731, and the National Institutes of Health grant R37GM065859 to B.L.B. J.E.S. is a Howard Hughes Medical Institute Fellow of the Jane Coffin Childs Memorial Fund for Medical Research. O.P.D. was supported by the NIGMS T32GM007388 grant. F.A.H. is funded by the Schmidt Science Fellowship. K.J.F. was supported by the Endowed Scholars Program at the University of Texas Southwestern Medical Center, a Searle Scholars award, and NIH grant 1DP2-AI154402. The content is solely the responsibility of the authors and does not necessarily represent the official views of the funders.


#### AUTHOR CONTRIBUTIONS

J.E.S., O.P.D., F.A.H., K.J.F., and B.L.B. conceptualized the project. J.E.S. and O.P.D. constructed strains. J.E.S and O.P.D. performed experiments. J.E.S., F.A.H., and K.J.F. performed bioinformatic analyses. J.E.S., O.P.D., F.A.H., K.J.F. and B.L.B. analyzed data. J.E.S., O.P.D., and B.L.B. designed experiments. J.E.S., O.P.D., and B.L.B. wrote the paper.

#### COMPETING INTERESTS

The authors declare no competing interests.

#### ADDITIONAL INFORMATION

### EXTENDED DATA

#### Extended Data Figure 1

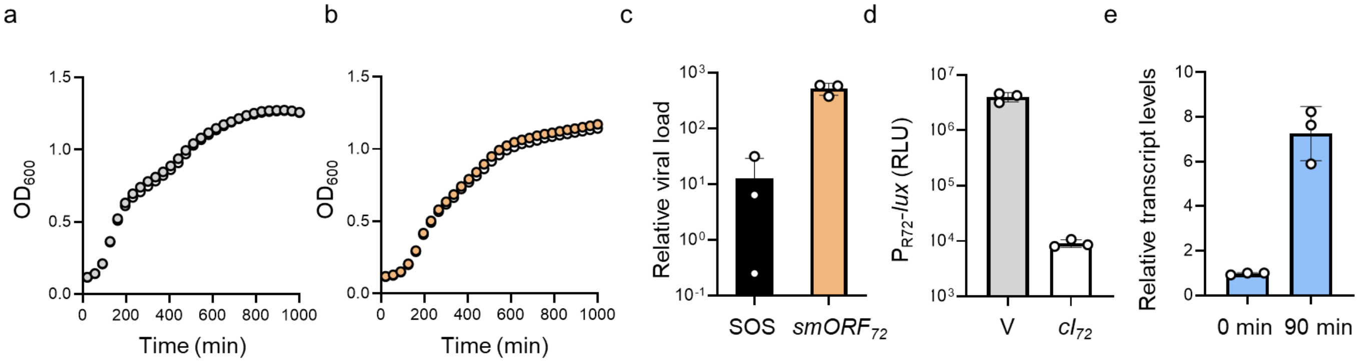


**Extended Data Figure 1. smORF$_{72}$-induced lysis of *Vibrio* 1F-97 is phage dependent and requires a TF-smORF module.**
**(a)** Growth of WT *Vibrio* 1F-97 in medium lacking or containing aTc (white and gray, respectively).
**(b)** Growth of *E. coli* carrying aTc-inducible *smORF$_{72}$* in medium lacking or containing aTc (white and orange, respectively).
**(c)** Detection of phage 72-specific particles in viral preparations obtained from culture fluids from *Vibrio* 1F-97 carrying aTc-inducible *smORF$_{72}$* that were grown in medium with ciprofloxacin or aTc to induce SOS and smORF$_{72}$ production, respectively. Relative viral load is the amount of *cI$_{72}$* DNA in the induced samples relative to that in an uninduced sample as judged by qPCR.
**(d)** Expression of plasmid-borne P$_{R72}$-*lux* in *E. coli* containing an empty vector (V) or the phage *cI$_{72}$* gene.
**(e)** Relative expression of *smORF$_{72}$* as judged by RT-qPCR in *Vibrio* 1F-97 carrying aTc-inducible TF$_{72}$ at 0 min and 90 min after induction with aTc. Relative transcript levels are the amount of phage *smORF$_{72}$* RNA relative to the amount of *rpoB* RNA, normalized to T=0 min.
Data are represented as means ± std with *n*=3 biological replicates (a, b, d) and as means ± std with *n*=3 biological replicates and *n*=4 technical replicates (c, e). RLU as in Figure 1d (d). aTc; 50 ng mL$^{-1}$ (a, b, c, e), ciprofloxacin; 500 ng mL$^{-1}$ (c).

### Extended Data Figure 2

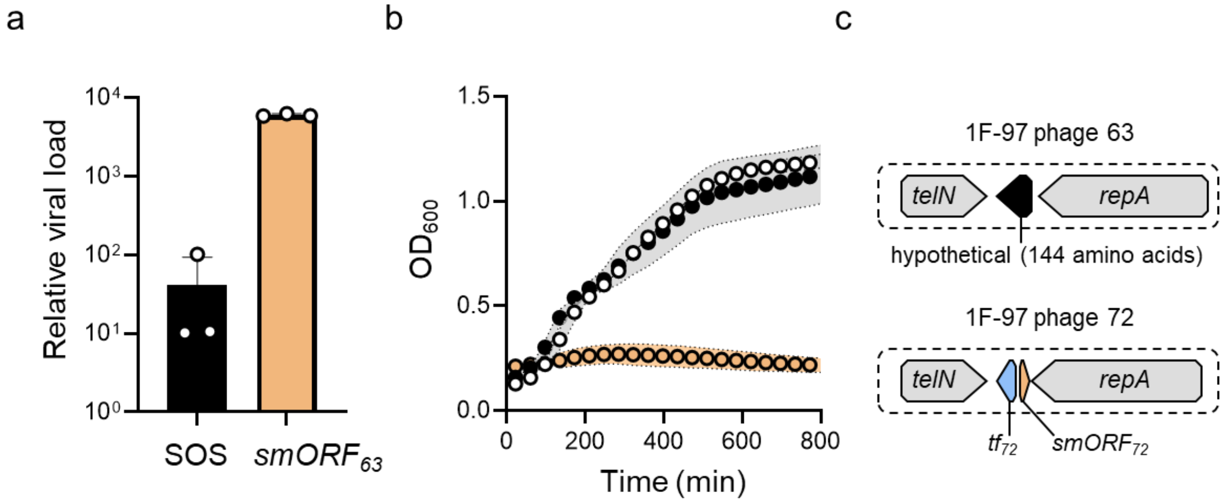


**Extended Data Figure 2. Genes encoding a TF-smORF module in phage 63 are located outside of the *repA*-*telN* region on the genome and the module induces lysis of *Vibrio* 1F-97.**

**(a)** Detection of phage 63-specific particles in viral preparations of culture fluids from *Vibrio* 1F-97 carrying aTc-inducible $smORF_{63}$ that were grown in medium with ciprofloxacin or aTc to induce SOS and $\text{smORF}_{63}$ production, respectively. Relative viral load is the amount of $cI_{63}$ in the induced samples relative to an uninduced sample as judged by qPCR.

**(b)** Growth of *Vibrio* 1F-97 carrying either a plasmid containing the intervening gene between *repA and telN* from phage 63 under an aTc-inducible promoter (black), aTc-inducible $smORF_{72}$ on a plasmid (orange), or no plasmid (white). All media contained aTc.

**(c)** Genetic organization of genes encoded between *repA* and *telN* in phage 63 and phage 72.

Data are represented as means ± std with *n*=3 biological replicates (b) and as means ± std with *n*=3 biological replicates and *n*=4 technical replicates (a). aTc; 50 ng $mL^{-1}$ (a, b), ciprofloxacin; 500 ng $mL^{-1}$ (a).

### Extended Data Figure 3

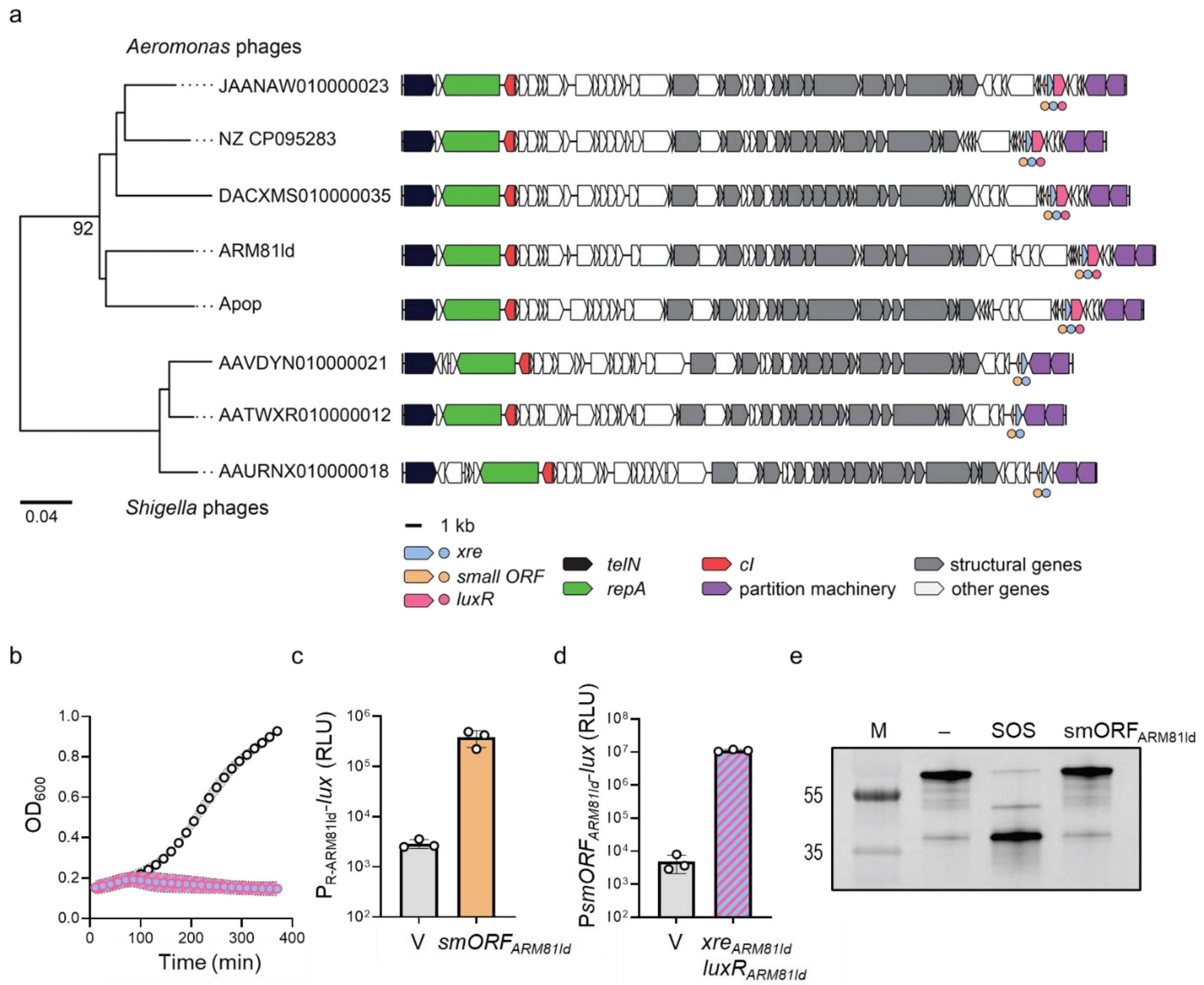


**Extended Data Figure 3. Comparison of linear plasmid-like phages in *Shigella* and *Aeromonas* reveals similar locations of genes encoding TF-smORF regulatory modules and similar functions among module components.**

**(a)** Phylogenetic tree (left) of 5 *Aeromonas* phages encoding *xre-luxR* genes and 3 *Shigella* phages encoding *xre*, with the genome organizations of each sequence depicted at right. Genes are colored by annotation as noted in the key below the figure. Circles denote the locations of relevant features that are common among the 8 phages (*xre* (blue) and *smORF* (orange)) or exclusive to the 5 *Aeromonas* phages (*luxR* (pink)). *Shigella* phages are labeled with their corresponding NCBI accession numbers. Numbers above branches are pseudo-bootstrap support values from 100 replications.

**(b)** Growth of *Aeromonas* sp. ARM81 harboring aTc-inducible $xre_{ARM81ld}$-$luxR_{ARM81ld}$ in the presence and absence of aTc (blue/pink and white, respectively). All media contained C4-HSL.

**(c)** $P_{R\text{-}ARM81ld}$-*lux* expression from *E. coli* carrying an empty vector (V) or aTc-inducible $smORF_{ARM81ld}$ in medium containing aTc. The $P_{R\text{-}ARM81ld}$-*lux* plasmid carries two copies of $cI_{ARM81ld}$ (see Methods) for native repression of reporter expression.

**(d)** $PsmORF_{ARM81ld}$-*lux* expression in *E. coli* carrying an empty vector (V) or aTc-inducible $xre_{ARM81ld}$-$luxR_{ARM81ld}$ in medium containing aTc and C4-HSL.

**(e)** SDS-PAGE in-gel labeling of the ARM81ld repressor (HALO-$cI_{ARM81ld}$) produced in *E. coli* carrying aTc-inducible $smORF_{ARM81ld}$. The treatments -, SOS, and $\mathrm{smORF}_{\mathrm{ARM81ld}}$ refer to water, ciprofloxacin, and aTc, respectively. M as in Figure 1b.

Data are represented as means ± std with *n*=3 biological replicates (b, c, d). RLU as in Figure 1d (c, d). aTc; 50 ng $mL^{-1}$ (c, d, e), aTc; 5 ng $mL^{-1}$ (b), ciprofloxacin; 500 ng $mL^{-1}$ (e), C4-HSL; 10 µM (b, d).
